# Integrative Genomic Analyses Identify *COL21A1* and *ENPEP-FGF5* Regulatory Pathways for Blood Pressure Variation in East Asians

**DOI:** 10.64898/2026.05.14.725285

**Authors:** Zhi Chng Lau, Xuling Chang, Kar Seng Sim, Hao Wu, Arshia Naaz, Umamaheswari Muniasamy, Chiea Chuen Khor, Woon-Puay Koh, Vitaly A. Sorokin, Rajkumar Dorajoo

**Author notes:** Corresponding authors. Rajkumar Dorajoo: Laboratory of Metabolic Disease and Aging Genomics, Genome Institute of Singapore (GIS), Agency for Science, Technology and Research (A*STAR), 60 Biopolis Street, Genome, Singapore 138672, Singapore.

## Abstract

**Background:** Hypertension is a highly heritable cardiovascular disorder and a major determinant of cardiometabolic disease, including diabetes. However, the regulatory genes and tissue-specific mechanisms underlying blood pressure variations remain incompletely understood.

**Methods:** Leveraging a well-characterized prospective population-based cohort comprised of 27,308 participants from the Singapore Chinese Health Study (SCHS), we evaluated genome-wide genetic associations for five blood pressure traits: hypertension status, systolic blood pressure, diastolic blood pressure, mean arterial pressure (MAP) and pulse pressure (PP). Additionally, we conducted a transcriptome-wide association study (TWAS), integrating gene expression data from 49 tissues, followed by colocalization and fine-mapping to prioritize regulatory genes. Association of identified variants with incident diabetes was additionally evaluated in the longitudinal data.

**Results:** We validated 10 blood pressure loci (*P* between 1.64 x 10^-20^ – 4.10 x 10^-8^) and identified an East-Asian specific splice donor variant at the *COL21A1* gene associated with PP (rs149344559, *P* = 6.78 × 10^-10^). Integrative analyses prioritized *FGF5* in kidney cortex and *ENPEP* in pituitary tissue as candidate regulatory genes. The blood pressure-lowering allele at *ENPEP* (T allele, rs1879056) was associated with reduced risk of incident diabetes. Mediation analysis demonstrated that approximately 21% of the genetic association with diabetes was mediated through MAP (P_indirect-effect_ = 2 x 10^-16^).

**Conclusion:** This study refines genetic predispositions for blood pressure among East-Asians. We further delineate tissue-specific regulatory pathways underlying blood pressure variations and identify *ENPEP*-mediated dysfunctions linking blood pressure genetics to diabetes risk, underscoring integrated disease mechanisms.

## Introduction

Hypertension, characterized by persistently elevated systemic arterial blood pressure, affects over one billion people worldwide and is a baseline risk factor for various adverse cardiovascular outcomes [1, 2, 3]. Genetic factors account for approximately 30% to 60% of inter-individual variation in blood pressure [4].

To date, over a hundred genomic loci have been reported for systolic blood pressure (SBP), diastolic blood pressure (DBP) and hypertension in different populations by genome-wide association studies (GWAS) [4, 5, 6]. However, these identified genetic factors explain typically less than 10% of the phenotypic variance in blood pressure [3, 5]. Moreover, most GWAS risk loci are located in the non-coding regions of the genome and the target genes and tissues through which they influence blood pressure remain incompletely understood.

A major hypothesis is that these non-coding variants influence blood pressure through tissue-specific regulation of gene expression. Hence, integrating gene expression data may overcome the challenges in GWAS to investigate underlying biological mechanisms and understand how these genetic variants may functionally influence blood pressure regulation *in vivo* [7, 8]. Recently, transcriptome-wide association studies (TWAS) have emerged as a gene-based analytical framework for complex traits analysis. By integrating GWAS summary data with expression quantitative trait loci (eQTLs) reference panels, TWAS enables the identification of gene-trait associations in a tissue-specific and directionally informative manner [8].

In this study, we performed GWAS on five blood pressure traits: hypertension status, SBP, DBP, mean arterial pressure (MAP) and pulse pressure (PP), in approximately 26,000 adults from the Singapore Chinese Health Study (SCHS). SCHS a population-based prospective cohort that has been actively followed up for over 20 years in Singapore. Subsequently, we performed TWAS to identify tissue-specific dysregulated genes associated with blood pressure traits. For the prioritized variants, we further evaluated their associations with incident diabetes to investigate potential downstream clinical implications. We hypothesized that integrating GWAS with tissue-specific transcriptomic regulation in a longitudinally followed East-Asian cohort would identify regulatory genes linking blood pressure genetics to cardiometabolic disease outcomes.

## Method

### Study population

SCHS is a long-term population-based prospective cohort study investigated on dietary, genetic and environmental determinants of cancer and other chronic diseases in Singapore [9, 10, 11]. Between April 1993 and December 1998, 63,257 Singaporean Chinese participants aged 45 to 74 years old from two major Chinese dialect groups in Singapore, the Hokkien and the Cantonese, were recruited. At recruitment, participants underwent in-person interviews using structured questionnaires to collect information on demographics, anthropometric measurement, lifetime use of tobacco, alcohol consumption, usual diet, physical activity, menstrual/reproductive history (women only), and medical history. After recruitment, consenting survivors were re-contacted for Follow-up 1 (1999-2004) and Follow-up 2 interviews (2006-2010) where information on lifestyle factors and disease status was updated. In addition, biospecimens (blood or buccal cells, and urine samples) were collected from a total of 32,575 consenting participants between 1999 and 2004.

At baseline, diabetes status was assessed based on self-reported physician diagnosis, and individuals with a documented history of diagnosed diabetes were excluded from the analysis. Incident diabetes cases were defined as participants who reported developing diabetes in subsequent follow-ups [12]. Participants were asked: “Have you been told by a doctor that you have diabetes (high blood sugar)?” If the answer was “yes”, participants were also asked for the age at which they were first diagnosed. The accuracy of self-reported diabetes has been estimated to be 98.8% in the SCHS [13].

All study procedures were approved by the Institutional Review Board (IRB) at the National University of Singapore (LB-16-241). Written informed consent was obtained from all participants.

### Blood pressure traits

SBP and DBP were measured three times within a 3-minute interval, and the mean of three readings rounded up to the nearest integer was used for analysis. As previously reported, SBP and DBP were adjusted for anti-hypertensive medication usage, where an addition of 15mm/Hg and 10mm/Hg were added to the original SBP and DBP respectively if the individual was on medication [14]. MAP was calculated as (2 x DBP + SBP)/3 and PP was calculated as SBP – DBP. Both MAP and PP were derived using medication-adjusted SBP_adjusted_ and DBP_adjusted_ values. Individuals who were on anti-hypertensive medication, SBP_adjusted_ exceeding 140mm/Hg or DBP_adjusted_ exceeding 90mm/Hg, were defined as those with hypertension.

### Genome-wide association study

A total of 27,308 DNA samples from SCHS participants were whole genome genotyped using the Illumina Global Screening Array (GSA). This included 18,114 samples genotyped on v1.0 and 9,194 samples genotyped on v2.0. Comprehensive information regarding genotyping procedures and quality control (QC) protocols have been described previously [9].

To augment the dataset, additional autosomal single nucleotide polymorphisms (SNPs) were imputed using local population-specific reference panels obtained from the SG10K whole-genome sequencing initiative (SG10K Health) [15, 16] on the Research Assets Provisioning and Tracking Online Repository (RAPTOR). All SNP alleles were standardized to the forward strand and mapped to the GRCh38 reference genome. Minimac4 (version 1.0.0) was employed to impute variants in the SCHS study, utilizing a set of 9,770 local Singaporean population samples as reference panels, as derived from the SG10K study [16]. High-quality imputed common SNPs were defined as those with minor allele frequency (MAF) ≥ 0.01 and those with an impute r² value ≥ 0.30 (N = 7,264,695).

### Statistical analysis for GWAS

Quantitative blood pressure traits were normalized to z-scores and analyzed using linear models implemented in Plink2 [17], adjusting for relevant covariates (age, sex and the top three principal components of population stratification (PCs 1-3)). Hypertension status (cross-sectional binary trait) was analyzed using the Scalable and Accurate Implementation of GEneralized mixed model (SAIGE v1.1.5) [18], which applies saddlepoint approximation (SPA) to account for case-control imbalance. Effect sizes estimation was provided through Firth’s bias-reduced logistic regression.

The associations of SNPs with the five blood pressure traits evaluated in the study, and the statistical models and software used for individual analyses were detailed in Supplementary Table 1. We used a P-value cut-off of *P* < 5 × 10^-8^ in the study to identify genome-wide significant associations. All significant hits as well as variants in linkage disequilibrium (LD; r² > 0.2, East Asian reference from the 1000 Genome reference population) were characterized using FUMA GWAS, GWAS catalog and OpenTargets [19, 20, 21, 22] to evaluate their consistency with previous GWAS sentinel SNP findings.

### Transcriptome-wide association study

Pre-computed gene expression reference weights were downloaded from the FUnctional Summary-based ImputatiON (FUSION) website (http://gusevlab.org/projects/fusion/), derived from the latest release version of Genotype-Tissue Expression (GTEx) v8 multi-tissue expression dataset. Only genes with nominally significant *cis*-SNP heritability (*P* < 0.05) within ±500 kb were included for predictive modeling. The sample sizes used to compute gene expression reference weights and the transcript number of genes covered were listed in Supplementary Table 2. Detail description of tissue collection, biospecimen processing and QC procedures on measured gene expression and genotype information for GTEx v8 reference panels are described in [23, 24].

FUSION evaluates multiple predictive models, including best linear unbiased predictor (BLUP), Bayesian sparse linear mixed models (BSLMM), Least Absolute Shrinkage and Selection Operator (LASSO), elastic net and Top1 models. For each gene-tissue pair, fivefold cross-validation was performed to compute out-of-sample *R*², and the model with the highest significant predictive performance was selected.

TWAS association statistics were computed using Z_TWAS_=w+Z/(w[Lw]1/2), where w denotes the SNP expression weights, Z denotes GWAS Z-scores, and L denotes the LD correlation matrix [8]. Statistical significance was evaluated using a two-tailed test under the standard normal distribution. The TWAS significant association was determined using Bonferroni-correction for 28,021 genes at *P_TWAS_* < 1.78 x 10^-6^. Detail information about the principle of FUSION, statistical model, and Z_TWAS_ calculation were described in the original paper [8]. LD reference panels were derived from East Asian samples in the 1000 Genomes Project to match cohort ancestry.

Conditional joint analyses were performed to distinguish jointly significant genes from marginally associated genes in regions with multiple TWAS signals. We also quantified the proportion of GWAS association explained by predicted gene expression [25, 26].

### Colocalization and fine-mapping analyses

Colocalization analyses were performed using COLOC software [27] to assess whether GWAS and eQTL signals shared a common causal variant. Posterior probabilities were computed for five hypotheses (PP0 – PP4), with strong evidence for colocalization defined as PP4 > 0.8.

Statistical fine-mapping was conducted using FOCUS (fine-mapping of causal gene sets), which models predicted expression correlations induced by LD and TWAS gene expression reference weights [28]. Posterior inclusion probability (PIP) was calculated for each gene, and 90% credible gene sets were constructed. Genes with PIP > 0.5 were considered candidate regulatory genes.

### Association and mediation analyses for incident diabetes

We evaluated the association using the most significant GWAS SNP at each TWAS-prioritized locus with incident diabetes in the SCHS cohort. Regression models were adjusted for age, sex and population substructure (PCs 1-3).

For variants significantly associated with incident diabetes, mediation analyses were performed using the Structural Equation Modeling (SEM) module in STATA (ver 15) to estimate the total effect, direct effect and indirect effect through mediating variable of the SNPs on those phenotypes, which showed significant associations [29]. Subsequently, the proportion of SNP effects mediated was calculated as the proportion of indirect effect over the total effect. All mediation analysis was adjusted for age, sex, and population substructure (PCs 1-3).

## Results

### Common variant genetic associations

Figure 1 provides a visual representation of the overall study design. Descriptive data for the demographics of the SCHS cohort is presented in Supplementary Table 3. The mean ± standard deviation (SD) for age of study participants at recruitment was 54.75 ± 7.23 years and 43.8% (*N* =10,342) were men.

**Figure 1:**
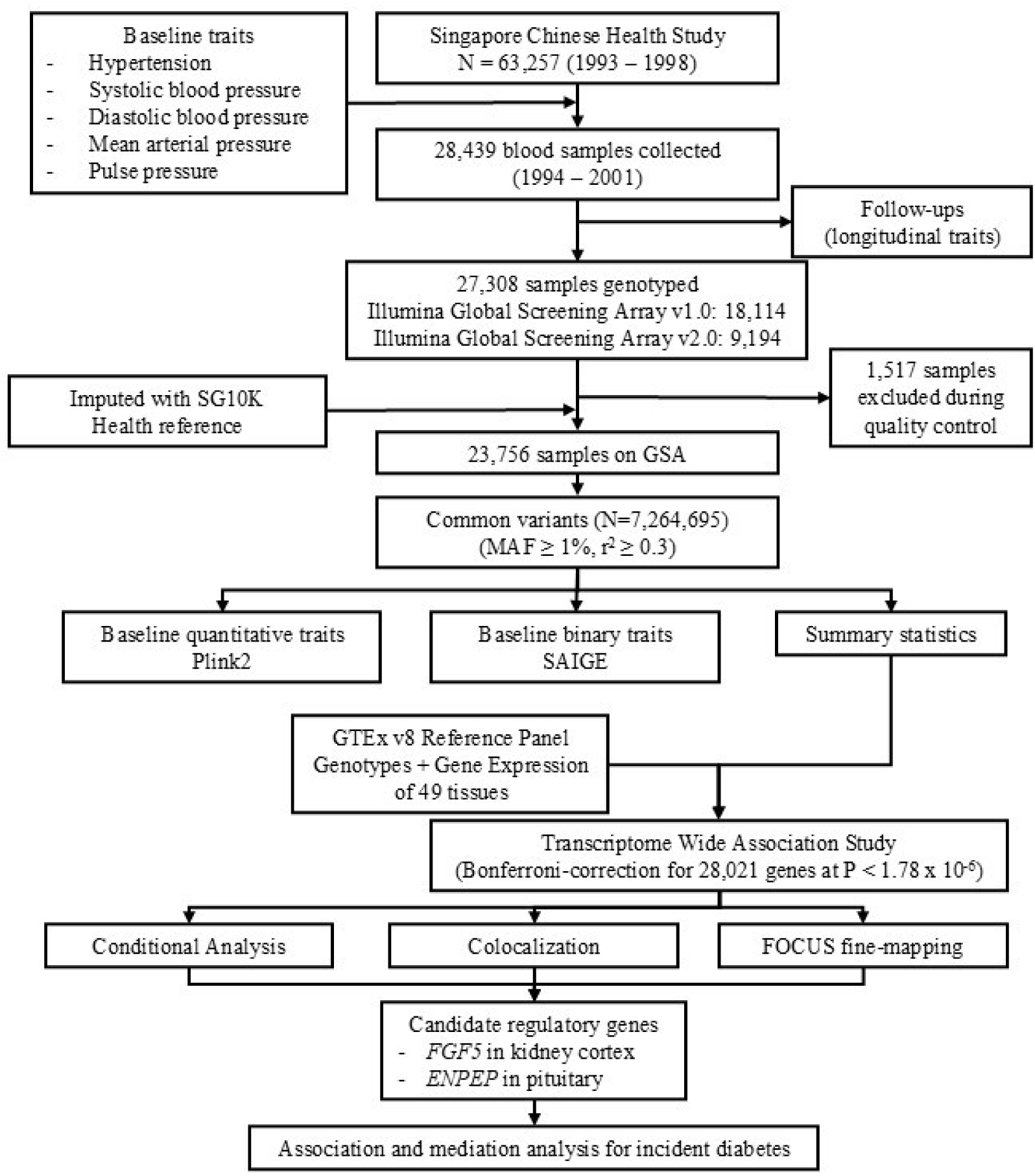
Study design of GWAS and TWAS for blood pressure traits. The SCHS cohort recruited approximately 63,000 Singaporean Chinese participants from two major Chinese dialect groups in Singapore, the Hokkien and the Cantonese. Information on blood pressure traits was measured and collected at recruitment. Approximately half of the cohort donated blood or buccal samples for DNA extraction and whole genome genotyped. Association of high quality imputed common SNPs with blood pressure traits were evaluated. TWAS was performed by integrating the predictive models from the Genotype-Tissue Expression (GTEx) reference panel with the GWAS summary data for blood pressure traits. Conditional, colocalization, and fine-mapping analyses were performed to determine the candidate regulatory genes associated with blood pressure. Finally, prioritised genes were evaluated for associations with incident diabetes in SCHS.

Genetic association analyses were performed using variants with high quality imputation scores (MAF ≥ 1%, r^2^ ≥ 0.3). We identified 10 robust genetic associations for blood pressure traits and hypertension status at previously reported loci that were beyond the genome-wide threshold of significance (Supplementary Table 4) [30, 31, 32]. These replicated genetic signals in the SCHS dataset were largely the same reported sentinel SNPs or in strong linkage disequilibrium (LD; r^2^ > 0.6, 1000G CHS reference panel) with the sentinel SNPs reported for the corresponding trait.

### Genetic associations with pulse pressure

We identified one additional locus that was associated with PP (rs149344559, *P* = 6.78 x 10^-10^, Table 1). Rs149344559 is a splice donor variant at the *COL21A1* gene with a high Combined Annotation-Dependent Depletion (CADD) score of 34 and likely an East-Asian specific variant [monomorphic or very rare (MAF < 0.1%) in non-EAS reference populations].

**Table 1:**
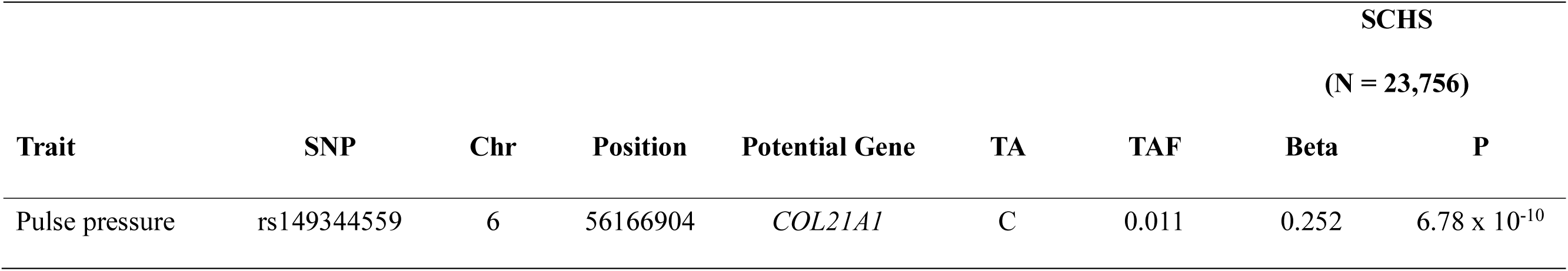
Summary statistics of genetic associations identified in the study. We identified one locus that was associated with pulse pressure. TA: test allele; TAF: test allele frequency.

Although *COL21A1* is a known locus for PP, our top hit (rs149344559), was not in LD with other sentinel SNPs previously reported for PP at this locus [33]. In a conditional probability analysis, the rs149344559-PP association remained statistically significant after adjusting for the previous hits in this locus, i.e. rs12201429 [33] (*P*_condition_ = 6.94 × 10^-10^) and rs1474698 [34] (*P*_condition_ = 2.32 × 10^-8^) (Supplementary Table 5). Additional evaluations of rs149344559 in the Taiwan Biobank pheweb (https://taiwanview.twbiobank.org.tw/pheweb.php) [31], showed robust associations with PP (*P* = 1.40 × 10^-8^), confirming this East-Asian specific association.

### TWAS analysis for blood pressure traits

We performed TWAS by integrating genome-wide summary statistics for blood pressure traits from 23,756 participants in the SCHS with gene expression reference weights across 49 tissues from the GTEx v8 reference panel. For significant TWAS associations across the 49 tissues (defined as Bonferroni-correction for 28,021 genes at *P* < 1.78 x 10^-6^), we identified 31 associations (representing 10 unique genes) for SBP, 4 associations (representing 2 unique genes) for DBP, 19 associations (representing 8 unique genes) for MAP, 22 associations (representing 5 unique genes) for PP, and 6 associations (representing 4 unique genes) for hypertension status (Supplementary Table 6).

Of these, five genes including *COL21A1*, *CUBN*, *GML*, *PRDM8* and *WNT2B* exhibited significant associations in only one tissue, whereas others showed multi-tissue associations. The most significant gene across all blood pressure traits was *FGF5* in kidney cortex. The direction of effect was consistent across traits, with Z_TWAS_ values ranging from 5.56 to 9.87, (*P_TWAS_* range from 5.13 x 10^-23^ to 2.67 x 10^-8^).

Conditional analyses were performed to evaluate independence of TWAS-significant genes within shared loci. The genes that were jointly and marginally significant were summarized in Supplementary Table 7 and 8. We further quantified the proportion of GWAS variance explained by TWAS-derived gene-expression associations. The variance explained ranged from 54.4% to 100% (Supplementary Table 8). Notably, *LSP1* and *PRR33* in cell cultured fibroblasts fully accounted for the corresponding GWAS variance for both SBP and MAP. Similarly, *ENPEP* expression in pituitary and spleen fully explained the GWAS variance in MAP. These findings indicate that BP-associated variants at these loci are likely mediated through genetically regulated gene expression.

### Colocalization and FOCUS fine-mapping analyses

Colocalization analyses were conducted to determine whether TWAS and GWAS signals shared common causal variants. We observed strong evidence for colocalization (PP4 > 0.8) for 25/31 SBP associations at eight unique genes, 3/4 DBP associations at two unique genes, 17/19 MAP associations at seven unique genes, 21/22 PP associations at four unique genes and all significant associations for hypertension status (4 unique genes) (Supplementary Table 6).

We then performed statistical fine-mapping using FOCUS to estimate the PIP for each gene being the likely regulatory gene at each locus. Across all blood pressure traits, *FGF5* in kidney cortex showed the strongest effects (PIP range from 0.997 to 1.00), supported by strong colocalization signals (PP4 > 0.8) (Table 2, Supplementary Table 9). Additionally, *ENPEP* in pituitary was identified as a likely regulatory gene for MAP (PIP = 0.55).

**Table 2:** FOCUS fine-mapping results for TWAS associations that also showed strong evidence of colocalization (PP4 > 0.8). FOCUS estimates the posterior inclusion probability of each gene being regulatory within a region of association, using the sum of posterior probabilities to define the default 90% credible set, a set of genes likely to contain the regulatory gene. Genes with PIP values > 0.5 were candidate regulatory genes. SBP: Systolic blood pressure; DBP: Diastolic blood pressure; MAP: Mean arterial pressure; PP: pulse pressure; FOCUS PIP: Posterior Inclusion Probability estimated by FOCUS. FOCUS region: Genomic region FOCUS fine-mapping results correspond to.

| **Trait** | **TWAS Location** | **Tissue** | **Gene** | **TWAS Z** | **TWAS P** | **FOCUS PIP** | **FOCUS region** |
| --- | --- | --- | --- | --- | --- | --- | --- |
| Hypertension | chr4:80266598-80266599 | Kidney cortex | *FGF5* | 9.88 | $5.13 \times 10^{-23}$ | 1 | 4:80070271-4:81201964 |
| SBP | chr4:80266598-80266599 | Kidney cortex | *FGF5* | 8.24 | $1.78 \times 10^{-16}$ | 1 | 4:80070271-4:81201964 |
| DBP | chr4:80266598-80266599 | Kidney cortex | *FGF5* | 9.00 | $2.36 \times 10^{-19}$ | 1 | 4:80070271-4:81201964 |
| MAP | chr4:80266598-80266599 | Kidney cortex | *FGF5* | 9.00 | $2.22 \times 10^{-19}$ | 1 | 4:80070271-4:81201964 |
| | chr4:110365732-110365733 | Pituitary | *ENPEP* | -5.01 | $5.53 \times 10^{-7}$ | 0.55 | 4:110336296-4:112947451 |
| PP | chr4:80266598-80266599 | Kidney cortex | *FGF5* | 5.56 | $2.67 \times 10^{-8}$ | 0.997 | 4:80070271-4:81201964 |

### Association of prioritised genes with incident diabetes

Variants at *FGF5* and *ENPEP* showed strong associations across blood pressure traits (*P* between 2.15 x 10^-4^ to 2.22 x 10^-19^, Supplementary Table 10). We next performed association analysis to evaluate whether the TWAS-prioritized candidate genes were associated with incident diabetes in our longitudinal cohort.

At *ENPEP*, the blood pressure-lowering allele (T allele, rs1879056) is associated with reduced risks of incident diabetes (HR = 0.92, 95% CI: 0.87 - 0.97, *P* = 2.81 x 10^-3^). TWAS results further indicated that decreased *ENPEP* expression in pituitary tissue was associated with lower MAP, supporting a regulatory role for *ENPEP* in blood pressure biology. Mediation analysis demonstrated that MAP significantly mediated the association between rs1879056 and incident diabetes, with 21.3% of the effect explained through MAP (*P*_indirect-effect_ = 2 x 10^-16^, Figure 2). Notably, these findings provide convergent genetic evidence across GWAS, TWAS and mediation analyses, implicating *ENPEP* as a regulatory node linking blood pressure and metabolic disease.

**Figure 2:**
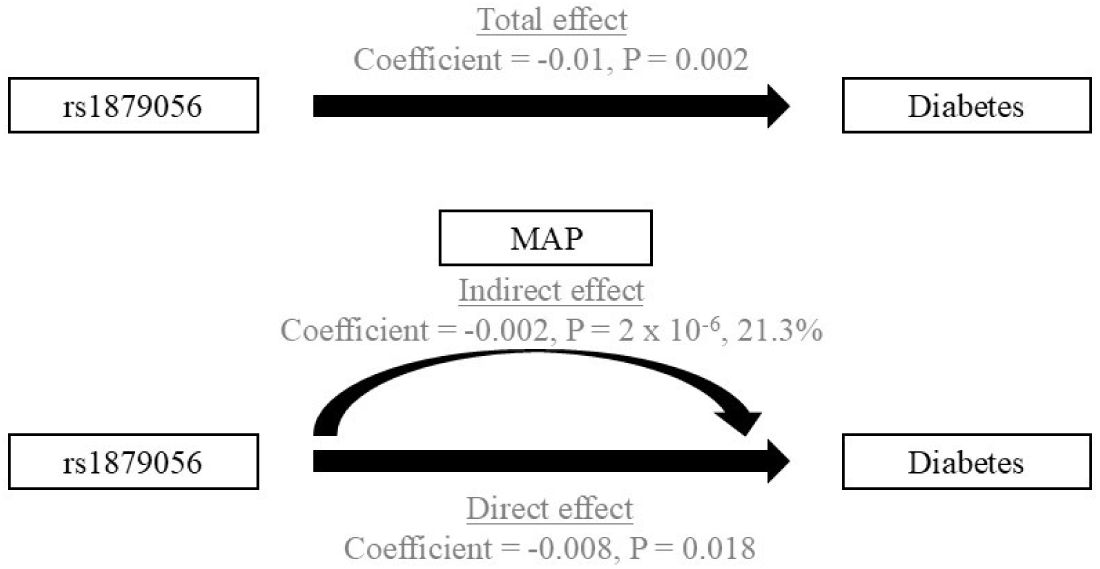
*ENPEP* variant associated with incident diabetes. Genetic predisposition of *ENPEP* (rs1879056) was associated with reduced incident diabetes, and mediation analysis demonstrated approximately 21% of the genetic association with diabetes was mediated through mean arterial pressure (MAP).

## Discussion

In this study, we systematically performed genetic associations and TWAS across five blood pressure traits and followed-up observed associations with longitudinal data on incident diabetes in a well-curated population dataset of Singaporean Chinese adults that has been actively followed-up for over 20 years. Our study corroborated several genetic associations, highlighting their transferability to the East-Asian ancestry population as well as identified an additional SNP association that were relevant to the East-Asian population.

Rare variant at the *COL21A1* has been previously associated with PP in individuals of European ancestry [35, 36]. *COL21A1* encodes the collagen alpha1 chain precursor of type XXI collagen, a fibril-associated collagen with interrupted helices (FACIT) that is localized to the extracellular matrix (ECM) and contributes to the structural organization of type I collagen-containing tissues [37]. As a minor collagen, *COL21A1* is thought to modulate collagen fibril assembly and ECM integrity, thereby influencing vascular structure and mechanical properties. Notably, patients with severe coronary artery disease, defined by high SYNTAX scores, exhibit elevated type I collagen deposition in the aortic wall compared to those with lower scores, highlighting the role of ECM remodeling in vascular pathology [38]. In this context, our findings of an independent East-Asian specific splice donor suggest a potential role of vascular ECM remodeling in PP.

TWAS and fine-mapping analyses consistently prioritized *FGF5* in kidney cortex as the top candidate regulatory gene across all blood pressure traits (PIP range from 0.997 to 1.00). *FGF5* (Fibroblast Growth Factor 5) has been repeatedly implicated as a hypertension susceptibility gene, with genetic variants and circulating expression levels positively correlated with blood pressure [39, 40, 41]. However, most prior studies relied on peripheral blood expression profiles. Our findings extend this body of work by demonstrating that genetically regulated expression of *FGF5* in kidney cortex (an organ of key relevance to blood pressure regulation) likely mediates the observed GWAS associations. Furthermore, CRISPR-based genome-editing studies have shown altered gene expression following deletion of the *FGF5* regulatory region [40, 41], suggesting that targeted modulation of this locus may provide a tractable value for functional validation and therapeutic exploration.

*ENPEP* emerged as a tissue-specific candidate gene in pituitary tissue. *ENPEP* encodes glutamyl aminopeptidase, a mammalian type II integral membrane protein with an extracellular zinc-binding domain, is closely involved in the well-known catabolic pathway of the renin-angiotensin system (RAS), facilitating conversion of angiotensin II to angiotensin III. Angiotensin III remains biologically active at AT1 receptors and is particularly relevant in central and pituitary RAS signaling [35, 42]. Immunodetection studies have identified a local RAS system in the pituitary [43], and *ENPEP* knockout mouse models develop hypertension [44], supporting its functional role in blood pressure regulation.

A central finding of our study is that blood pressure partially mediates the association between *ENPEP* variants and incident diabetes. A significant 21% of the effect of rs1879056 on diabetes risk was mediated through MAP, implicating a biologically meaningful pathway. Hypertension and diabetes share bidirectional pathophysiological mechanisms such as the upregulation of RAS, microvascular dysfunction, oxidative stress, inflammation, and activation of immune system [45, 46]. *ENPEP* expression in pituitary tissue may attenuate central RAS activity, leading to lower MAP and, consequently, reduced hemodynamic and microvascular stress that contributes to diabetes development. These results position *ENPEP* as a potential mechanistic link between hypertension and metabolic disease, highlighting central RAS modulation as a plausible pathway connecting blood pressure regulation to glycemic outcomes. However, further functional validation through *in-vitro* and *in-vivo* studies are necessary to clarify tissue-specific effects and potential systemic implications.

An advantage of the study was the well characterized longitudinal collection of data that enabled systematic identification and follow-up assessments of risk variants. Integration of GWAS with tissue-resolved eQTL reference panels allowed for prioritization of regulatory genes and provided mechanistic insight into the biological pathways related to blood pressure. However, several limitations should be considered. Data from the GTEx study are largely obtained from non-diseases individuals and may also not fully capture disease-relevant gene expression signatures and or context specific regulatory effects pertinent to hypertension and diabetes. TWAS also relies on genetically predicted gene expressions and does not directly measure transcriptional activity. Nevertheless, the integration of multiple complementary analyses in our study that included TWAS, colocalization and fine-mapping provides convergent evidence that strengthens gene prioritization. Future larger scale studies incorporating larger disease-relevant transcriptomic datasets alongside other genomic data such as proteomics and metabolomics, would be important to improve resolution of tissue-specific regulatory effects and refine causal inferences.

In conclusion, our study highlights the value of performing genetic evaluations of blood pressure traits and integrating GWAS with tissue-specific gene expression datasets. We identified ancestry-specific variation at *COL21A1*, confirmed kidney cortex *FGF5* as a robust regulator of blood pressure, and highlighted pituitary *ENPEP* in linking blood pressure regulation to downstream metabolic disease risk. These findings highlight the role of renal and central pathways in blood pressure control and provide important insights into the genetic architecture of hypertension among East-Asian populations.

## Authors’ contributions

Zhi Chng Lau: Conceptualization, Methodology, Validation, Formal analysis, Investigation, Visualization, Project administration, Writing – original draft, Writing – review & editing. Xuling Chang: Formal analysis, Investigation, Writing – original draft, Writing – review & editing. Kar Seng Sim: Methodology, Formal analysis, Investigation, Writing – original draft, Writing – review & editing. Arshia Naaz: Investigation, Writing - review & editing. Hao Wu: Formal analysis, Visualization, Writing – review & editing. Umamaheswari Muniasamy: Methodology, Writing – review & editing. Chiea Chuen Khor: Methodology, Writing – review & editing. Woon-Puay Koh: Supervision, Writing – original draft, Writing – review & editing. Vitaly A. Sorokin: Conceptualization, Supervision, Writing – original draft, Writing – review & editing. Rajkumar Dorajoo: Conceptualization, Formal analysis, Methodology, Validation, Supervision, Funding acquisition, Writing – original draft, Writing – review & editing.

## Acknowledgments

W-P.K. was supported by the National Medical Research Council, Singapore [CSA-SI (MOH-000434)]. The Singapore Chinese Health Study was supported by the US National Institutes of Health (NIH) (grants R01 CA144034 and UM1 CA182876), the Singapore National Medical Research Council (NMRC/CIRG/1456/2016) and the Singapore Strategic Cohorts Funding (P2022-02-01). We thank Siew-Hong Low of the National University of Singapore for supervising the fieldwork in the Singapore Chinese Health Study.

## Data Availability

All significant summary statistics in the study are appended in Supplementary 4 and 5. All genetic data for the SCHS are housed in the RAPTOR genomics repository (https://raptor.gisapps.org/). The data that support the findings of our study are available from the corresponding author of the study upon reasonable request.

## Supplementary table legends

Supplementary Table 1: Overview of the blood pressure-related traits investigated in the study

Supplementary Table 2: Characteristics of the Gene Expression Reference Weights

Supplementary Table 3: Demographics of the study population

Supplementary Table 4: Replication of known hits at genome-wide level of significance

Supplementary Table 5: Conditional analysis for sentinel SNPs in SCHS

Supplementary Table 6: Statistically significant transcriptome-wide associations

Supplementary Table 7: Conditional joint analysis results - jointly significant associations

Supplementary Table 8: Conditional joint analysis results - jointly and marginally significant associations per locus

Supplementary Table 9: FOCUS fine-mapping results for TWAS significant associations (PIP > 0.5) that has strong evidence of colocalization (PP4 > 0.8)

Supplementary Table 10: Associations between the SNPs for blood pressure traits and incident diabetes in SCHS

